# Resting After Learning Facilitates Memory Consolidation and Reverses Spatial Reorientation Impairments in ‘New Surroundings’ in 3xTg-AD Mice

**DOI:** 10.1101/2024.11.12.622722

**Authors:** A. C. Stimmell, L. J. Alday, J. Marquez Diaz, S. C. Moseley, S. D Cushing, E. M. Salvador, S. M. Ragsdale, A. A. Wilber

**Affiliations:** Department of Psychology, Florida State University, Tallahassee FL USA; Department of Neuroscience; Wallace H. Coulter Department of Biomedical Engineering, Georgia Institute of Technology and Emory University, Atlanta GA USA; Neuroscience GIPD, University of Arizona, Tuscon AZ USA

## Abstract

Sleep is an essential component of productive memory consolidation and waste clearance, including pathology associated with Alzheimer’s disease (AD). Facilitation of sleep decreases Aβ and tau accumulation and is important for the consolidation of spatial memories. We previously found that 6-month female 3xTg-AD mice were impaired at spatial reorientation. Given the association between sleep and AD, we assessed the impact of added rest on impaired spatial reorientation that we previously observed. We randomly assigned 3xTg-AD mice to a rest (n=7; 50 min pre- & post-task induced rest) or a non-rest group (n=7; mice remained in the home cage pre- & post-task). Mice in both groups were compared to non-Tg, age-matched, non-rest controls (n=6). To confirm that our sleep condition induced sleep, we performed the same experiment with rest sessions for both 3xTg-AD and non-Tg mice (n=6/group) implanted with recording electrodes to capture local field potentials (LFPs), which were used to classify sleep states. Markers of pathology were also assessed in the parietal-hippocampal network, where we previously showed pTau positive cell density predicted spatial reorientation ability (pTau, 6E10, M78, and M22). However, we found that 3xTg-AD rest mice were not impaired at spatial reorientation compared to non-Tg mice and performed better than 3xTg-AD non-rest mice (replicating our previous work). This recovered behavior persisted despite no change in the density of pathology positive cells. Thus, improving sleep in early stages of AD pathology offers a promising approach for facilitating memory consolidation and improving cognition.

## 1 Introduction

Alzheimer’s disease (AD) is a debilitating neurodegenerative disorder characterized by amyloid beta (Aβ) and Tau protein aggregation (Braak and Braak, 1991; Mitchell et al., 2002), in addition to other pathological processes (Hansson et al., 2006; Ragsdale et al., 2024). In AD, pathological processes develop and progress significantly, well before cognitive symptoms manifest (McDade and Bateman, 2017). However, getting lost, particularly in new surroundings, is an early cognitive impairment in humans that develop AD (Allison et al., 2016; Henderson et al., 1989).These spatial symptoms can help to push the diagnostic timeline forward. Recent work, including our own, suggests that this early impairment could represent a failure to use distal cues to reorient in space (Stimmell et al., 2019), impaired consolidation of memories during sleep (Cushing et al., 2020), or likely both. Sleep disruption is a risk factor for AD (Ju et al., 2017; Mander et al., 2017; Shi et al., 2018; Winsky-Sommerer et al., 2019). The relationship between AD and sleep appears to be bidirectional, with AD pathology exacerbating sleep-wake cycle disturbances and impaired sleep furthering accumulation of Aβ (Ju et al., 2014; Ragsdale et al., 2024). Sleep also has homeostatic roles, such as promoting synaptic weakening of neurons and decreasing cellular stress, which allows restoration of energy reserves in the brain, and is necessary due to the high energy cost during wakefulness (Tononi and Cirelli, 2014).

In addition to the homeostatic role of sleep, it also plays a critical role in memory formation. During sleep, behaviorally relevant neural activity replays in the hippocampus in coordination with the neocortex, which facilitates memory consolidation (Euston et al., 2007; Maingret et al., 2016; Qin et al., 1997; Wilber et al., 2017; Wilson and McNaughton, 1994). We previously observed that female 3xTg-AD mice were impaired at a behavioral paradigm that required them to repeatedly reorient in space using distal cues (Stimmell et al., 2019). Consistent with a role of sleep in memory facilitation, subsequent work in our lab found that impaired hippocampal-cortical coupling during sleep may at least partially explain the impaired spatial reorientation we observed (Cushing et al., 2020).

Another important function of sleep is to help restore the brain by clearing waste metabolites that may be neurotoxic (Van Egroo et al., 2019; Wang and Holtzman, 2020; Xie et al., 2013). This is believed to largely be accomplished by the glymphatic system, a macroscopic waste clearance system which uses AQP4 water channels on astrocyte end-feet to promote removal of soluble proteins from the central nervous system (CNS; Hablitz et al., 2020). The two hallmarks of AD, Aβ and tau pathologies, have been shown to fluctuate diurnally throughout the sleep-wake cycle, with increases during wakefulness and decreases during sleep (Lucey, 2020; Nebel et al., 2018; Wang and Holtzman, 2020). The glymphatic system is dependent on circadian timing and has been observed to be more effective during sleep (Hablitz et al., 2020). This could be responsible for the diurnal fluctuations (but see: Miao et al., 2024). Blood-brain barrier (BBB) based clearance of amyloid and tau aggregates also occurs during sleep and thus could also play a role in reductions of aggregates during sleep (Deane et al., 2008). In people with AD, an increase of Aβ in cerebrospinal fluid has been shown to emerge with the onset of sleep impairments, such as increased sleep fragmentation (Cushing et al., 2020; Holth et al., 2017; Ju et al., 2013; Sprecher et al., 2017; Varga et al., 2016). AD is associated with disrupted sleep, poor sleep quality, and reduced sleep duration, leading to cycle of an increased accumulation of Aβ and tau (Holth et al., 2017). More specifically, disrupted sleep and SWS deficits are associated with increased Aβ (Branger et al., 2016; Sprecher et al., 2017; Yun et al., 2017), possibly because the glymphatic system is not able to effectively clear pathology (Roh et al., 2012). Animal models of AD also show sleep alterations occurring with Aβ deposition (Colas et al., 2004; Cushing et al., 2020; Huitrón-Reséndiz et al., 2002; Jyoti et al., 2015; Roh et al., 2012; Schneider et al., 2014; Wisor et al., 2005). Decreased sleep has been shown to appear in some mouse models after the formation of Aβ plaques (Huitrón-Reséndiz et al., 2002; Roh et al., 2012; Van Erum et al., 2019), while an amyloidosis-only mouse model (APP/PSEN) showed decreased sleep before the formation of Aβ plaques (Jyoti et al., 2015). Regardless of which comes first (sleep deficits or amyloid and tau aggregation), the bidirectional relationship between sleep and clearance acts as a positive feedback loop, further contributing to the development of neuroinflammation and pathology, thereby advancing disease progression (Ju et al., 2017).

A great deal of research examining the relationship between sleep and AD has focused on improving 24h circadian sleep because a common disorder in AD patients is irregularity of the sleep-wake cycle, with reduced amplitude of circadian rhythmicity, phase delay, and sundowning, which has been reproduced in animal models by numerous studies, including our own (Beesley et al., 2020; Leng et al., 2019; Volicer et al., 2001). However, much less work, and none in rodents (to our knowledge), have examined the possibility of using rest sessions to improve cognitive symptoms associated with AD. We also focused on mice that had intracellular accumulation of Aβ and tau but no plaques or tangles, coinciding with the emergence of deficits on our sensitive cognitive test, unlike previous studies which have focused on timepoints after plaques and/or tangles have formed. To assess the impact of sleep on early AD with spatial navigation, we used the spatial reorientation task. This task requires mice to repeatedly reorient in space using distal cues, but we added both pre- and post-task sleep sessions. We focused primarily on 6-month female 3xTg-AD mice since this group was previously shown to have spatial reorientation learning and memory deficits (Stimmell et al., 2019). Additionally, 6-month female 3xTg-AD mice showed pTau positive cell density profile across a parietal-hippocampal network which predicted spatial reorientation ability (Stimmell et al., 2021). Here, we assess spatial reorientation as well as markers of AD pathology to test if improving sleep early in disease progression improves cognition and if such improvements are related to Aβ and tau accumulation.

## 2 Methods

### 2.1 Animals

We assessed the effects of 1h rest sessions on spatial reorientation ability in two experiments. In the first experiment, sleep was manipulated under conditions nearly identical to those in which we previously observed cognitive deficits in this spatial learning and memory task (with a small brain stimulation implant; Stimmell et al., 2019). In the second experiment, we directly assessed sleep with a larger recording array implant. Overall, for both experiments, 6-month female 3xTg-AD (n = 20) and age-matched non-Tg control mice from the same background strain (n = 12) were grouped housed (2-4/cage) in 12:12 h light/dark cycles until the beginning of the experiment. These mice underwent behavioral testing during the 12h light phase of the light/dark cycle in a dimly lit room. For Experiment 1, mice (3xTg-AD, n = 14; non-Tg, n = 6) did not have a recording array implant to ensure that the effects we observed in Experiment 2 were not a consequence of the recording array. The 3xTg-AD mice in Experiment 1 were subdivided into no-sleep (n=7) and sleep (n= 7) groups. For Experiment 2, a subset of the mice (n = 6/genotype) had a recording array implant (some data from a different and later training phase was published previously; Cushing et al 2020), which allowed us to measure the quantity of slow-wave sleep (SWS) and rapid eye movement (REM) sleep. All experimental procedures were carried out in accordance with the NIH Guide for the Care and Use of Laboratory Animals and approved by the University of California, Irvine (n = 1 mouse) and Florida State University Animal Care and Use Committee (n = 31 mice).

### 2.2 Behavioral Assessment

The behavioral assessments for pre-training, stimulation parameters, spatial reorientation task, and probe testing were performed using the same as the methods as in Stimmell et al. (2019) so that two groups in this study were a replication of our previous work (3xTg-AD no-sleep mice and age-matched non-Tg controls).

### 2.3 Pre-training

Mice were water deprived to no less than 80% of their beginning body weight and given food *ad libitum*. They were then trained to shuttle to a black barrier at the end of a linear track and back for a water reward (*alternation training*). This barrier was positioned so that there was a black background behind it (from the view of the mouse) on the wall. This track was moved to different starting positions, so the length of the track varied (**Fig. 1**). The starting position was randomly selected from 9 possible start locations for both *alternation training* (spaced over a range of 56– 76 cm from the center of the goal zone). Note, all calculations were performed in pixel coordinates. The distances listed in cm throughout the paper were estimated for visualization purposes. After reaching criterion (either asymptote, ±6 total runs, 3 out of 4 days, or a total of 50 or more runs down the track and back), the mouse was scheduled for surgery to implant stimulating electrodes and continued with alternation training every other day until the day before surgery when water deprivation ceased.

**Figure 1.**
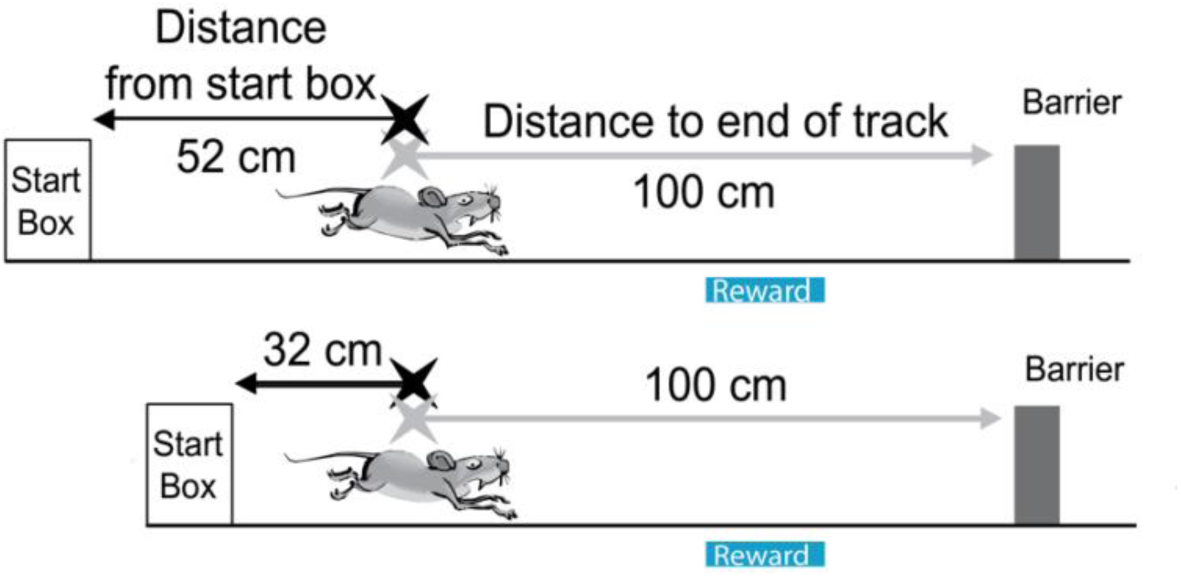
Spatial reorientation task. The maze consists of a linear track with a start box and an unmarked reward zone (blue bar). It is always fixed within the room; however, the start box and track move between trials. As they move down the track, the mice need to use stable external cues in the real to locate the reward zone and slow to obtain a brain stimulation reward. Adapted from Cushing et al., 2020 and Rosenzweig et al., 2003.

### 2.4 Stimulating Electrode Only Implantation

The surgical procedure for Experiment 1 was described previously (Stimmell et al., 2019). Once criteria were met on the alternation training task, mice underwent surgery to implant bilateral stimulating electrodes targeting the left and right medial forebrain bundle (MFB; 1.9 mm posterior to bregma, 0.8 mm lateral, 4.8 mm below dura).

### 2.5 Recording Array Implantation

The surgical procedure for the subset of animals with a recording array in Experiment 2 was described previously (Cushing et al., 2020). For this subset, two bipolar stimulating electrodes were implanted unilaterally, targeting the left MFB (1.9 mm & 1.4 mm posterior to bregma, ±0.8 mm lateral, 4.8 mm below dura). The implantation of these stimulating electrodes allowed for intracranial stimulation of the medial forebrain bundle. A 16-tetrode recording array (Chang et al., 2013) was implanted, targeting the parietal cortex and dorsal hippocampus (2.2 mm posterior to bregma, 2.0 mm lateral). Animals recovered for one week, during which tetrodes targeting the hippocampus were turned down approximately 124 µm daily for the first three days, and then 31 µm every other day, until the tetrodes reached hippocampus, as indicated by depth records, as well as the characteristic hippocampal local field potential (LFP). Data from tetrodes targeting the parietal cortex are not reported here. Tetrode locations were histologically verified *post-mortem*, based on the presence of damage tracks and/or an electrolytic lesion corresponding to the tetrode tip.

### 2.6 Stimulation Parameters

Following a 1-week recovery period, mice were placed in a custom 44 × 44 × 44 cm box with a circular nose poke port (Med Associates) mounted on the left half of one wall of the chamber. Using manual stimulations, mice were first shaped to approach the nose poke port, then to touch the port, and finally to poke their nose into the port to break an infrared beam. Once trained to nose poke, the animal automatically triggered each brain stimulation reward. Brain stimulation rewards lasting 500ms were delivered using a custom-written MATLAB (The MathWorks Inc.) program attached to a Stimulus Isolator (World Precision Instruments). Over the course of one week, settings were adjusted (171–201 Hz frequency, 30–70 μA current, electrode wire combinations) to achieve maximal response rate. No attempt was made to balance responding across genotype or age; however, responding (nose poke) rate was compared across genotype to ensure that differences in reward strength were not likely to contribute to the observed effects. The ideal measure of stimulation effectiveness is use of a running wheel to trigger brain stimulation because operant conditioning is less effective in mice (Carlezon and Chartoff, 2007). However, since our task requires mice to stop running for a reward, we did not want to confound the study by training mice to run for brain stimulation and then training them on the reverse (stopping for brain stimulation, i.e., reversal learning). Thus, we first assessed brain stimulation responding with an operant task, and this worked well for nearly all the mice in the study. For low responding mice, we assessed stimulation responsiveness with a task that required alternating for stimulations at either end of the track. This was assessed after spatial reorientation training, to prevent a confound of reversal learning. We then combined responses for stimulation across these two tasks, which produced a continuous distribution with all stim alternation values falling below operant training values.

### 2.7 Spatial Reorientation Training

Once mice were responding maximally, one additional refresher alternation training session was conducted, followed by spatial reorientation training on a mouse version of a task we published previously (Stimmell et al., 2021, 2019) and adapted from the Rosenzweig et al. (2003) task for rats. Throughout the remainder of training and testing, mice shuttled back and forth for a water reward delivered in the start box, which they consumed while the track was moved slowly and smoothly to the next randomly selected starting location. There were 9 possible start locations, spaced over a range of 56–76 cm from the center of the goal zone, decided using random lists generated by Random.org (random with repeats). Next, an unmarked reward zone was added to the task (28 cm from the barrier at the end of the track) that could automatically trigger a single brain stimulation reward lasting 500 ms. This reward was only delivered if the mouse remained in the zone for a sufficient period of time before progressing to the end of the track (**Fig. 1**). This zone was fixed in relation to the room and cues positioned around the periphery of the room and was marked in camera coordinates. Thus, there were no visible markings in the room or on the track indicating the location of the reward zone. Further, since the track was moved to new positions following each trial, any olfactory or other cues on the track could not signal the location of the reward zone. Thus, the reward zone occurred at a variety of physical locations on the track, but always in a fixed location within the room. If the mouse remained in the zone for the duration of a delay period (starting at 0.5 s), a brain stimulation reward was delivered. The delay was fixed for a given day and was increased by 0.5 s (up to 2.5 s) each time the mouse met a criterion of similar percent correct (±15%), 3 out of 4 days. Custom MATLAB software was used for timestamping the key events of each trial (start, reward zone, and end of track), automatically delivering the brain stimulation reward after the *reward delay*, and generating a record of integrated events and the tracked position of the mouse for offline analysis. We have previously made the software for running this experiment freely available (Brea Guerrero et al., 2023).

There were pre- and post-task sleep sessions before and after each daily session, excluding pretraining. These sessions were included while determining optimal stimulation parameters, and throughout the spatial reorientation task and probe testing. Each day consisted of a 50 min pre-task sleep session, followed by 20 min of task, and then a 50 min post-task sleep session (**Fig. 2b)**. Sleep was induced by placing mice in a 17 x 16.5 x 16 cm box while connected to the recording system (as in Cushing et al., 2020). As we have done previously (e.g., Stimmell et al. 2019), we assessed the average stimulation response rate to verify that a particular group was not responding better than the other. We found no significant effects of group in Experiment 1 (3xTg-AD sleep vs. non-Tg vs. 3xTg-AD no-sleep) on response rate (F_(2, 17)_ = 1.65, *p* = 0.22). Though unlikely to influence the results given that responding did not differ across groups, it is possible that higher settings were needed to produce this similar responding in these mice. Therefore, we also performed the same comparison for the two brain stimulation parameters that we adjusted: frequency and current. Non-recording array mice showed no variation across genotype for frequency (F_(2, 17)_ = 0.43, *p* = 0.66) or current (F_(2, 17)_ = 0.85, *p* = 0.44). Similarly, the two recording array mice groups from Experiment 2 (3xTg-AD vs. non-Tg) showed no differences across genotype in response rate (t_(10)_ = 0.69, *p* = 0.51). There was also no difference in frequency or currency across genotype (t_(10)_ = 0.52, *p* = 0.61; t_(10)_ = 0.15, *p* = 0.88; as reported in Cushing et al. 2020).

**Figure 2.**
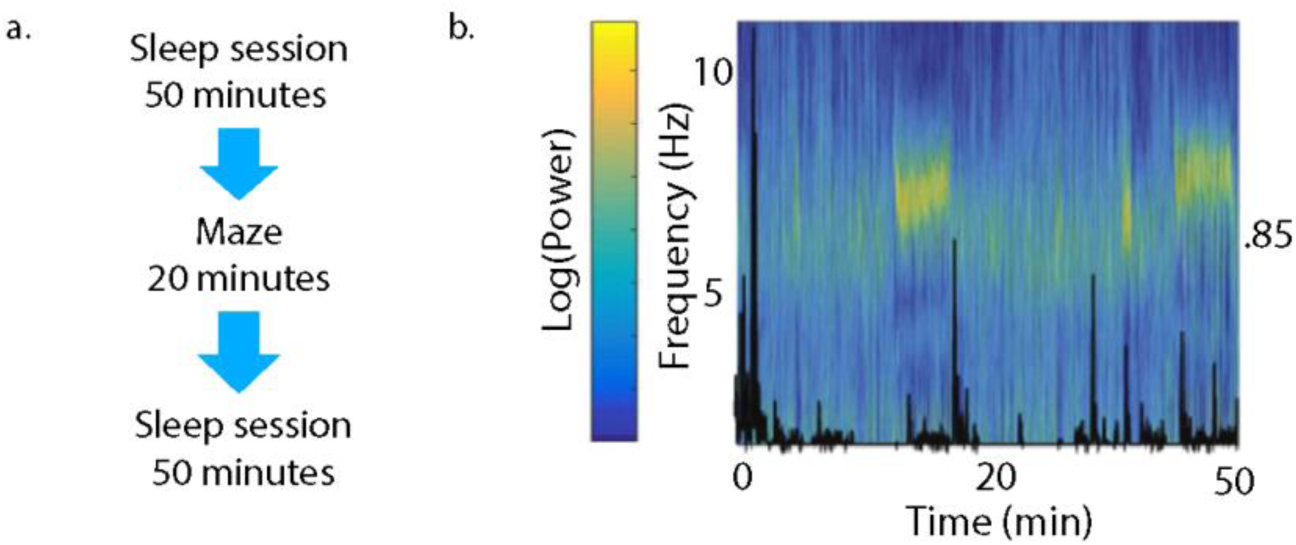
Differentiation of sleep states. **a.** Recording sessions for sleep mice started with a *pre-task* sleep session, followed by the task, then a *post-task* sleep session. Sleep sessions were 50 min each and task sessions lasted 20 min for each daily recording session. **b.** Time frequency power spectrum for LFP overlaid with movement velocity (black line) illustrating the contrast between slow-wave sleep (SWS; low velocity & low theta power), REM sleep (low velocity & high theta), and awake (high velocity & low theta; Buzsáki et al 2003, Grosmark et al 2012, Lansink et al 2008, Leung 1998, Mizuseki et al 2011, Mizuseki et al 2009, Sirota et al 2008). Figure adapted from Cushing et al., 2020.

### 2.8 Sleep Recording Procedures

As described in Cushing et al. (2020), an electrode interface board (EIB-72-QC-Small or EIB-36-16TT, Neuralynx) was attached with a custom adapter to the recording array utilizing independently drivable tetrodes (Chang et al., 2013) connected via a pair of unity-gain headstages (HS-36, Neuralynx) to the recording system (Digital Lynx SX, Neuralynx). Tetrodes were referenced to a tetrode wire in the corpus callosum and one wire from each tetrode was selected for LFP recording. Adjustments were made each day’s recording to allow for stabilization overnight. A continuous trace was collected for processing as LFP from one of the tetrode wires (bandpass-filtered 0.1 – 1000 Hz and digitized at 6400 Hz). Mouse position was tracked using a colored dome of reflective tape for the spatial reorientation task and online position information was used to trigger MFB stimulation rewards. Video-tracking data was collected at 30 Hz and co-registered with LFPs and event timestamps. LFP analyses were performed using custom-written MATLAB code or Freely Moving Animal (FMA) Toolbox (https://fmatoolbox.sourceforge.net/). The LFP signal was collected at 6400 Hz and subsequently resampled to 2000 Hz for further analysis, using the MATLAB *resample* function.

### 2.9 Sleep Quantification

First, still periods for both recording array mice and non-recording array sleep mice were extracted from the rest sessions as described previously (Cushing et al., 2020; Euston et al., 2007; Wilber et al., 2017). The raw position data from each video frame was smoothed by convolution of both x and y position data with a normalized Gaussian function (standard deviation of 120 video frames). After smoothing, the instantaneous velocity was found by taking the difference in position between successive video frames. An epoch during which the velocity was constantly below 0.78 pixels/s (∼0.19 cm/s) for more than 2 min was considered a stillness period. All analyses of rest sessions were limited to these stillness periods. Stillness data was also collected in an identical manner for Experiment 1. For Experiment 2 only, SWS and REM sleep were identified during these still periods using K-means clustering of the theta/delta power ration extracted from the CA1 pyramidal layer LFP recorded during the stillness periods, (**Fig. 2b**, Buzsáki et al., 2003; Girardeau et al., 2009; Grosmark et al., 2012; Lansink et al., 2008; Leung, 1998; Mizuseki et al., 2011; Sirota et al., 2008), and this approach has been validated (Costa-Miserachs et al., 2003).

### 2.10 Histology and Imaging

After the conclusion of the experiment, mice (3xTg-AD no-sleep, n = 7; 3xTg-AD sleep, n= 7) were euthanized by administering an intraperitoneal injection of Euthasol and transcardially perfused with 1X phosphate-buffered saline (PBS), followed by 4% paraformaldehyde (PFA) in 1X PBS. Whole heads were post-fixed in PFA for 24 h to identify the placement of simulating electrodes in the medial forebrain bundle and tetrode tracks when appropriate. After brain extraction, an additional 24 h post-fixation period was carried out, followed by cryoprotection of the brain in a 30% sucrose solution. Coronal frozen sections were cut at a thickness of 40 µm using a sliding microtome. Sections were split into 6 evenly spaced series for histological analysis. One series was processed for c-Fos for a subset of the data for a different larger study not reported here. Whole slides were imaged using a Zeiss Axioimager M2 with a 10x objective (∼100x magnification) and stitched together using Zeiss ZEN Blue software. Histology procedures were conducted as previously described in Stimmell et al. (2019), Cushing et al. (2020) and Stimmell et al. (2021). In summary:

#### 2.10.1 M22 and M78

Free-floating sections were blocked then incubated for two days in either primary antibody anti-MOC22 (monoclonal, rabbit, Abcam 205339) or anti-MOC78 (monoclonal, rabbit, Abcam 205341). Following a wash, sections were then incubated overnight in secondary antibody Anti-rabbit-alexa-488 (goat). Sections were stained for NeuN by utilizing fluorescently-labeled primary antibody anti-NeuN-Cy3 (polyclonal, rabbit, Millipore, ABN78C). Sections were mounted with DAPI-containing antifade mounting media, coverslipped, and imaged.

#### 2.10.2 Phosphorylated Tau

Free-floating sections were blocked then incubated overnight in primary antibodies anti-phosphorylated tau AT8 (monoclonal, mouse, MN1020) and anti-NeuN (polyclonal, rabbit, Millipore, ABN78). Following a wash, sections were then incubated 6h in secondary antibodies anti-mouse-alexa-488 and anti-rabbit-alexa-594. Sections were mounted with antifade mounting media, coverslipped, and imaged.

#### 2.10.3 6E10

Sections were mounted prior to incubation in 4% PFA for 4 min, followed by 70% formic acid for 2-4 min. Sections were then blocked and incubated for two days in primary antibodies anti-β-amyloid 1-16 (mouse, clone 6E10, Biolegend) and anti-NeuN (polyclonal, rabbit, Millipore, ABN78). Sections were washed, then incubated for 5-6 h in secondary antibodies anti-mouse-alexa-488 and anti-rabbit-alexa-594. Whole slides were rinsed, coverslipped with DAPI-containing antifade mounting media, and imaged.

### 2.11 Region of Interest Analyses

The density of positive cells for M22, M78, phosphorylated tau, and 6E10 intracellular pathology was assessed in four predefined regions of interest: retrosplenial cortex, CA1 field of the hippocampus, subiculum, and parietal cortex. These regions were further subdivided into dorsal (RSCd and SUBd) and ventral (RSCv and SUBv) parts for the retrosplenial cortex and subiculum. Dorsal CA1 (CA1d) encompassed all sections rostral to a point 2.55 mm posterior to bregma, while ventral CA1 (CA1v) included all sections caudal to that same point. Seven regions were examined in total. Using the Allen Brain Atlas, regions of interest (ROI) were manually drawn based on cytoarchitecture to locate regional boundaries. Automated cell counts were completed using Zeiss Zen 3.6 (blue edition). Images were taken with a 10x objective (approximately 100x magnification) using a Zeiss Axio Imager M2 microscope. Outlines were manually drawn around the ROI in a 1:12 evenly spaced series throughout the brain using manual selection tool in Zen. The experimenter remained blind to groups. The number of M22, M78, pTau, and 6E10 labeled cells was then expressed as the proportion per unit of area.

### 2.12 Statistics

Statistical comparisons for recording array mice (Experiment 1) were performed using two-way repeated measures ANOVAs (genotype by delay or genotype by position on the track) and unpaired t-tests (for two group comparisons or planned comparisons). A similar approach was used Experiment 2 groups (non-Tg mice, 3xTg-AD mice, and 3xTg-AD sleep mice). Statistical comparisons of histology data between groups were performed using two-way repeated measures ANOVAs (group x ROI), followed by post-testing corrected for repeated testing when appropriate (e.g., following interactions). For all statistical analyses, p<0.05 was considered significant. Statistical analyses were performed using Statview and R.

## 3. Results

### 3.1 6-month 3xTg-AD no-sleep mice are impaired at spatial reorientation compared to the 3xTg-AD sleep group

#### Experiment 1

Previously, we found that 6-month (but not 3-month) female 3xTg-AD mice were impaired at spatial reorientation learning and memory (Stimmell et al., 2019). Here we assessed spatial reorientation ability in 3xTg-AD sleep mice compared to non-Tg and 3xTg-AD no-sleep mice. The non-Tg and 3xTg-AD no-sleep mice do not undergo the pre- and post-task sleep sessions, making them a replication of our previous study. In **Figure 3a**, we isolated velocity data (Z-scored) for the first half of the reward zone. This measure allows for comparison of performance across *reward delays* (i.e., across the range of task difficulty; *0.5 – 2.5 s*) between groups. We found that 6-month 3xTg-AD sleep mice appear to perform similarly to non-Tg control mice, slowing more in the approach into the reward zone compared to 3xTg-AD mice, particularly in the *1 and 1.5 s reward delays*. Velocity varied significantly across groups (**Fig. 3b**; F_(2, 68)_ = 3.77, *p* < 0.05). However, velocity did not vary significantly across delays (F_(4, 68)_ = 1.71, *p* = 0.16), nor was there a significant interaction between delay and genotype (F_(8, 68)_ = 0.68, *p* = 0.70). Planned comparisons to investigate the cause of the group main effect revealed that 3xTg-AD no-sleep mice performed significantly worse than non-Tg mice (F_(1, 12)_ = 6.54, *p* < 0.05). 3xTg-AD no-sleep mice also performed significantly worse than 3xTg-AD sleep mice (F_(1, 12)_ = 5.03, *p* < 0.05). Non-Tg mice and 3xTg-AD sleep mice did not differ from each other (F_(1, 11)_ = 0.017, *p* = 0.90).

**Figure 3.**
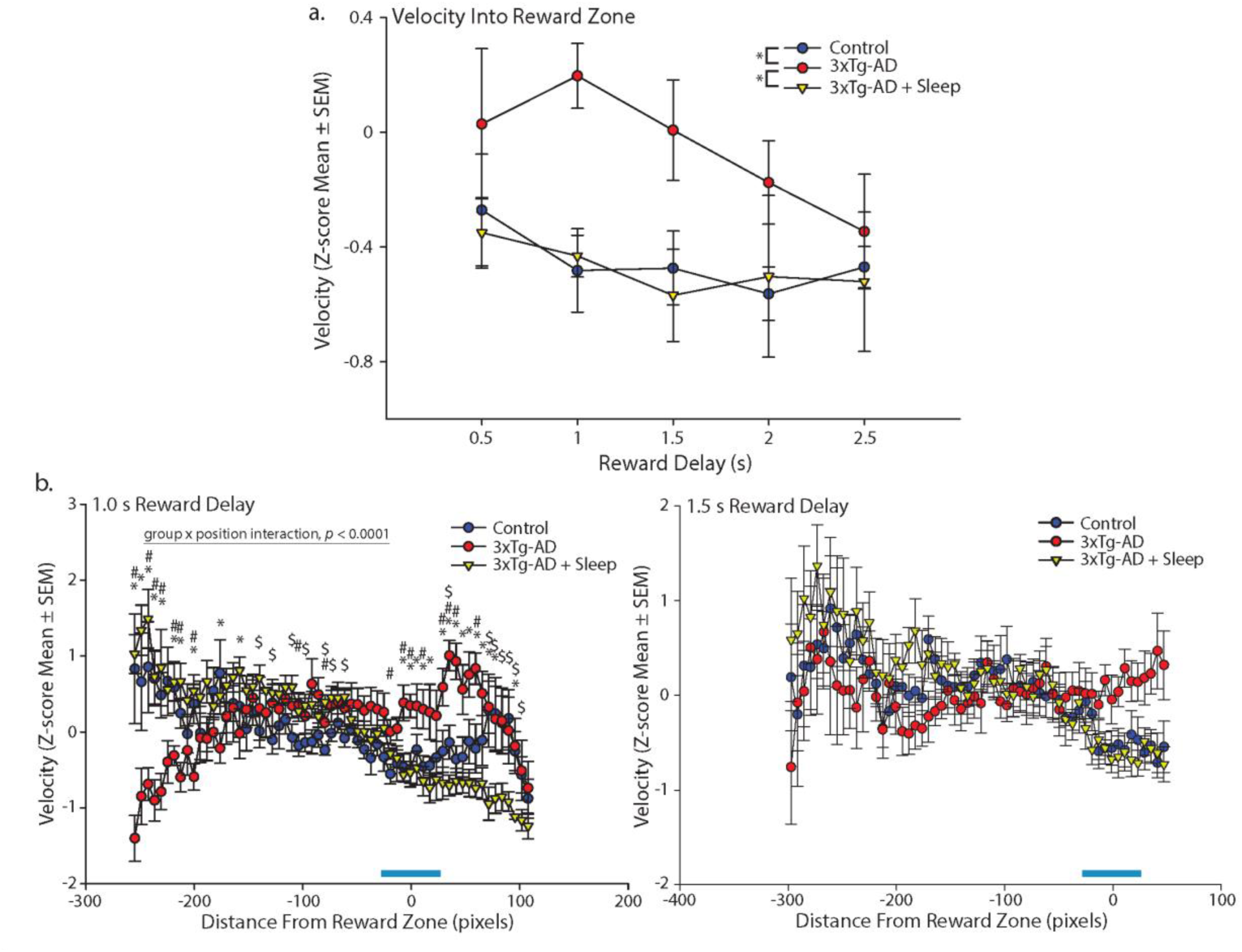
6-month 3xTg-AD female sleep mice are not impaired at spatial reorientation. **a**. Mean (±SEM) Z-scored velocity from the first half of the reward zone (left 1/2 of the blue bar in **b**) for each *reward delay* (*0.5-2.5 s*) for 6-month female non-Tg (blue) and 3xTg-AD (red) mice, as well as 6-month female 3xTg-AD sleep mice (yellow). 3xTg-AD sleep and non-Tg mice slow significantly more than 3xTg-AD mice with no sleep sessions (\**ps* < 0.05). **b.** *Left:* Mean (±SEM) Z-scored velocity plotted along the distance of the track during the *1 s reward delay* for 6-month female non-Tg mice, 6-month female 3xTg-AD mice, and 6-month female 3xTg-AD sleep mice. Both non-Tg and 3xTg-AD sleep mice slowed in the reward zone (blue bar). *Right:* Same as *left,* but for the *1.5 s reward delay*. 3xTg-AD sleep and non-Tg mice appear to slow more in the reward zone compared to controls, but not significantly. * 3xTg-AD vs. sleep mice *ps* <0.05; # 3xTg-AD vs. non-Tg mice *ps* <0.05; $ 3xTg-AD sleep vs. non-Tg mice *ps* < 0.05

Next, we assessed the velocity profile over the full length of the track (and not just the first ½ of the reward zone) during two intermediate *reward delays* with the greatest separation between groups on **Figure 3a**, the *1 and 1.5 s reward delays*. During the *1 s delay*, velocity did not vary significantly across groups (**Fig. 3b** *left*; F_(2, 1020)_ = 1.16, *p* = 0.34), but it did vary significantly across the length of the track (F_(60, 1020)_ = 3.63, *p* < 0.0001). The velocity profile also varied significantly as a function of group and position on the track (genotype by position interaction: F_(120, 1020)_ = 4.75, *p* < 0.0001). Planned comparisons showed that 6-month female 3xTg-AD sleep mice slowed more than 6-month female 3xTg-AD no-sleep mice for multiple locations on the track, close to and in the reward zone and after the reward zone (ts< -2.22, *ps* < 0.05; *s in **Fig. 3b** *left*), but towards the beginning of the track, an opposite effect was shown with 6-month female 3xTg-AD sleep mice running faster than 6-month female 3xTg-AD mice (ts > 2.22, *ps* < 0.05; *s in **Fig. 3b** *left*). Similarly, 6-month female non-Tg mice also slowed more than 6-month female 3xTg-AD no-sleep mice for multiple locations on the track prior to the reward zone, in the reward zone, and after the reward zone (ts > 2.22, *ps* < 0.05; #s in **Fig. 3b** *left*). An opposite effect also appears towards the beginning of the track, with 6-month female non-Tg mice running faster than 6-month female 3xTg-AD mice (ts < -2.58, *ps* < 0.03; #s in **Fig. 3b** *left*). In addition, 6-month female 3xTg-AD sleep mice ran more slowly than 6-month female non-Tg mice for multiple locations towards the end of the track and just prior to the reward zone (ts < -2.26; *ps* < 0.05, $s in **Fig. 3b** *left)*, but the opposite was seen in the middle of the track (ts > 2.32, *p* < 0.04; $s in **Fig. 3b** *left*). Together, this pattern of data suggests that 6-month female 3xTg-AD sleep mice are slowing in the reward zone and are performing similarly or slightly better than 6-month female non-Tg mice, while 6-month female 3xTg-AD no-sleep mice are not slowing as well for the reward zone. No-sleep 3xTg-AD mice are are also running more slowly over the earlier portions of the track, possibly indicating a lack of knowledge about the reward location.

During the *1.5 s delay*, velocity did not vary significantly across groups (**Fig. 3b** *right*; F_(2, 969)_ = 0.75, *p* = 0.49), but it did vary significantly across the length of the track (F_(57, 969)_ = 2.61, *p* < 0.0001). However, the velocity profile did not vary significantly as a function of group and position on the track (genotype by position interaction: F_(114, 969)_ = 1.16, *p* = 0.14). This suggests that all animals are adjusting their speed as a function of position on the track (potentially indicating some knowledge of the reward zone), but there are no significant differences between groups when considering data from the full length of the track.

### 3.2 6-month female 3xTg-AD sleep mice are still for a significant portion of the daily sleep sessions

We use a “boring” box to encourage mice to sleep, and our previous studies suggest that mice do spend significant time sleeping under these conditions (Cushing et al., 2020). However, those mice have a larger implant to allow for recording of brain activity patterns, which may further encourage immobility and sleep. Therefore, we next assessed stillness, defined as the amount of time they remained motionless for periods lasting at least 2 min, as a proxy for sleep in 3xTg-AD sleep mice (**Fig. 4**). We found that these mice were still (Cushing et al., 2020) approximately 40% of the time they were in the sleep box (i.e., about 40 min per day).

**Figure 4.**
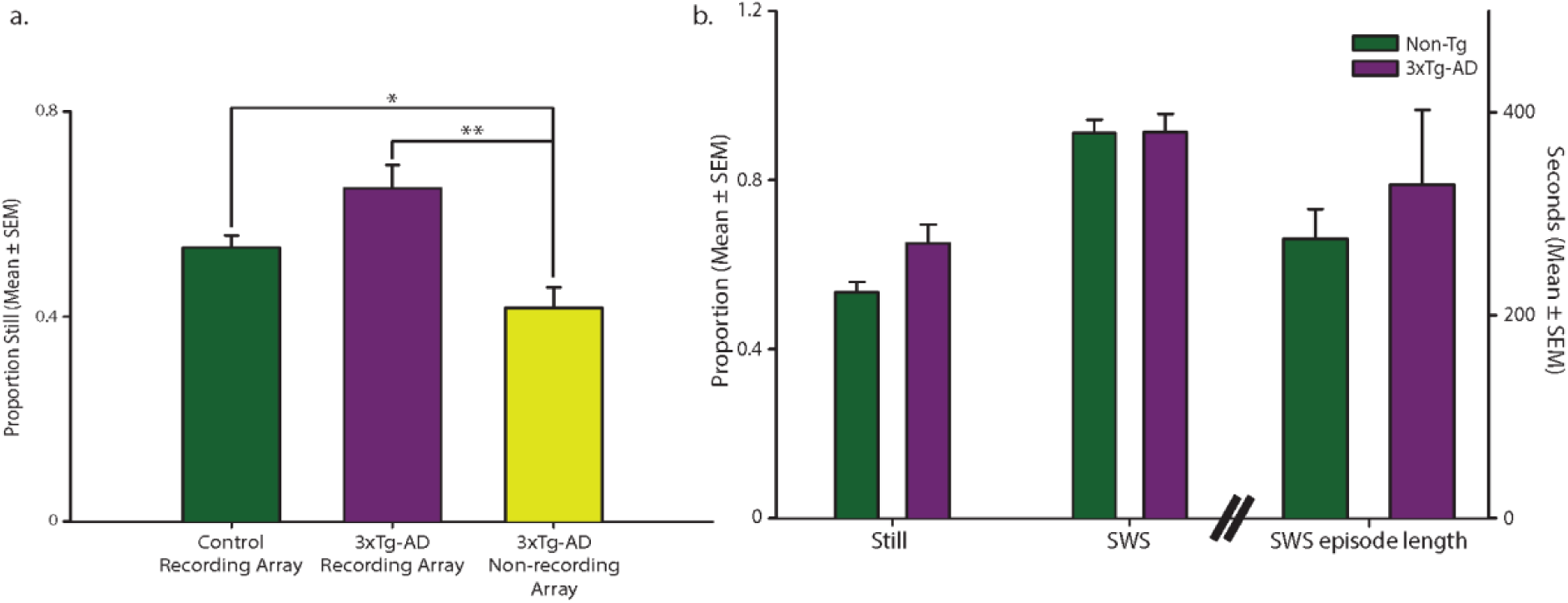
Sleep characteristics. **a.** 6-month 3xTg-AD sleep mice without a recording array spend less time still during sleep sessions. Mean (±SEM) proportion of stillness for non-Tg mice with a recording array (green), 3xTg-AD mice with a recording array (purple), and 3xTg-AD mice without a recording array (yellow). 3xTg-AD mice without a recording array spend significantly less time still during sleep sessions than both non-Tg and 3xTg-AD mice with a recording array. **b.** Sleep in 6-month female 3xTg-AD mice is similar to non-Tg recording array mice. Mean (±SEM) proportion of stillness, slow-wave sleep (SWS), and SWS episode length for non-Tg (green) and 3xTg-AD (purple) mice.

Furthermore, there was no significant difference in the proportion of stillness during pre-task and post-task sleep sessions (F_(1, 12)_ = 2.04, *p* = 0.18). There was also no significant effect of *reward delay* (F_(4, 48)_ = 1.80, *p* = 0.15) and no significant interaction between the sleep session and *reward delay* (F_(4, 48)_ = 1.02, *p* = 0.41) on sleep duration, suggesting that sleep did not change for the more difficult phases of the task. These data show that the mice were still for the same proportions during both sleep sessions and likely spent a significant amount of time sleeping during these rest sessions. One hour may be a critical time point for sleep dependent memory consolidation (Bourtchouladze et al., 1998). In the last ten minutes of both sleep sessions, approaching one hour, we found that these mice were still for approximately 80% of the last ten minutes at the 1.0s and 1.5s reward delays (*1.0s* Average = 0.81, SEM = 0.04; *1.5s* Average = 0.84, SEM = 0.03).

### 3.3 6-month 3xTg-AD sleep mice spend less time still than 3xTg-AD and Non-Tg mice with a recording array

Next, we compared stillness in mice with recording arrays to 3xTg-AD sleep mice (**Fig. 4a**). Stillness varied significantly across groups (F_(2, 14)_ = 9.22, *p* < 0.01). Planned comparisons between each group showed that 3xTg-AD sleep mice were still significantly less than both non-Tg (t_(10)_ = 2.25, *p* < 0.05) and 3xTg-AD recording array mice (t_(10)_ = 3.79, *p* < 0.01). Stillness did not vary significantly between non-Tg and 3xTg-AD recording array mice (t_(8)_ = 2.26, *p* = 0.05). These data suggest that the larger recording array leads to more sleep in 6-month mice, but that even the lower amount of sleep induced in 3xTg-AD sleep mice without a recording array is sufficient to reverse impairments in spatial reorientation learning and memory.

### 3.4 6-month 3xTg-AD sleep mice do not differ in sleep characteristics from Non-Tg mice

Finally, we assessed the average proportion of time spent in SWS (vs. REM sleep) during the pre- and post-task sleep sessions (**Fig 4b**). Non-Tg and 3xTg-AD recording array mice spent a similar proportion of time in SWS (t_(8)_ = 0.04, *p* = 0.97). Lastly, we assessed the average length of time for each SWS episode, and again, 3xTg-AD mice did not significantly differ from non-Tg controls (t_(8)_ = 0.67, *p* = 0.52). While no significant differences were observed, there was a trend toward an increased proportion of the sleep sessions spent still and an increased average length of SWS episodes for 3xTg-AD mice compared to non-Tg mice, consistent with our previous observations that these sleep characteristics are increased when these mice are learning a new task (Cushing et al., 2020). Thus, the sleep benefit for 3xTg-AD mice is not due to differences in sleep characteristics between 3xTg-AD mice and controls.

### 3.4 6-month female 3xTg-AD sleep mice with a recording array are not impaired at spatial reorientation learning and memory

#### Experiment 2

As a next step in assessing spatial reorientation ability in 3xTg-AD sleep mice, we compared 3xTg-AD sleep mice to non-Tg sleep controls with a recording array. Note that non-Tg mice in Experiment 1 did not undergo sleep sessions. Once again, we isolated Z-scored velocity data for the first half of the reward zone (**Fig. 5a**). We found that velocity for 6-month female mice varied significantly across delays (**Fig. 5a**; F_(4, 32)_ = 5.63, *p* < 0.05), indicating that task performance varied as difficulty increased. However, velocity did not vary significantly across genotype (F_(1, 32)_ = 1.48, *p* = 0.26), and there was no significant interaction between delay and genotype (F_(4, 32)_ = 1.71, *p* = 0.17). The 3xTg-AD group with sleep performed similarly to the non-Tg controls with sleep. This suggests that adding sleep sessions helped reverse the impairment previously observed in 3xTg-AD mice.

**Figure 5.**
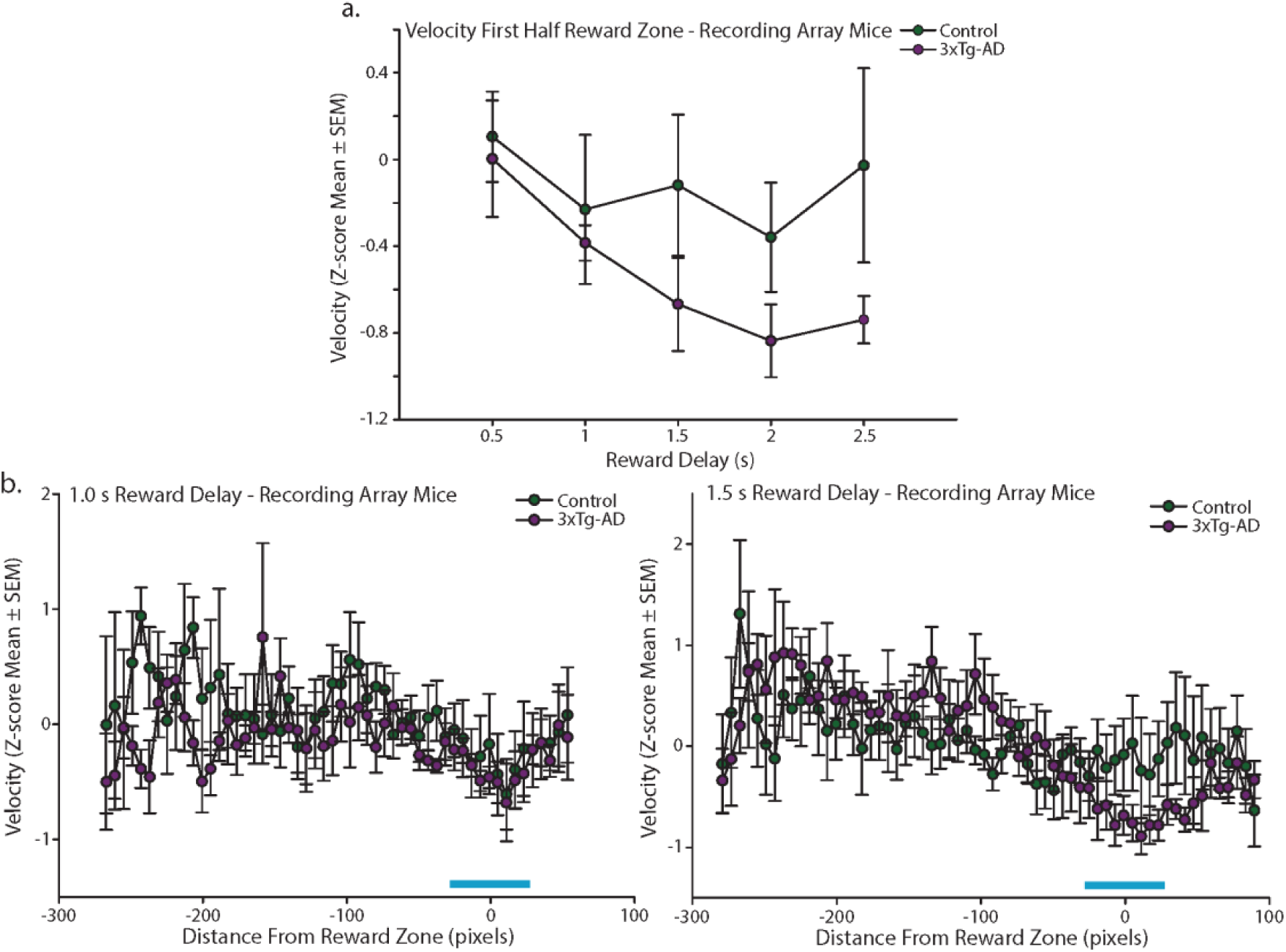
6-month 3xTg-AD female recording array mice are not impaired at spatial reorientation. **a.** Mean (±SEM) Z-scored velocity from the first half of the reward zone (blue bar in **b**) for each reward delay (*0.5-2.5 s*) for 6-month female non-Tg (green) and 3xTg-AD (purple) hyperdrive mice. 3xTg-AD recording array sleep mice did not perform significantly different than non-Tg recording array sleep mice. **b.** *Left:* Mean (±SEM) Z-scored velocity plotted for position on the track during the *1 s reward delay* for 6-month female non-Tg and 3xTg-AD recording array sleep mice. Both non-Tg and 3xTg-AD recording array sleep mice slowed into the reward zone (blue bar). *Right:* Same as *Left*, but for the *1.5 s reward delay*. 3xTg-AD recording array sleep mice did not significantly differ from sleep controls.

Next, we examined the mean Z-scored velocities for the full length of the track individually for two *reward delays* where we had observed group differences in Experiment 1 (the *1 and 1.5 s delays*; **Fig. 3b** *right*). During the *1 s delay*, velocity did not vary significantly across the length of the track (**Fig. 5b** *left*; F_(53, 424)_ = 0.98, *p* = 0.52), nor did it vary across genotype (F_(1, 424)_ = 4.20, *p* = 0.07). The velocity profile also did not vary significantly as a function of genotype and position in the room (genotype by position interaction: F_(53, 424)_ = 0.61, *p* = 0.99). During the *1.5 s delay*, velocity varied significantly across the length of the track (**Fig. 5b** *right*; F_(61, 488)_ = 2.29, *p* < 0.0001), but it did not vary across genotype (F_(1, 488)_ = 1.47×10^-4^, *p* = 0.99). The velocity profile also did not vary significantly as a function of genotype and position in the room (genotype by position interaction: F_(61, 488)_ = 0.95, *p* = 0.59). This pattern of data suggests that 6-month non-Tg and 3xTg-AD female sleep mice with a recording array mice were also performing similarly.

### 3.5 6-month Non-Tg mice perform similarly with or without sleep

To assess if sleep was benefitting 6-month female recording array control mice, we assessed their velocity compared to 6-month female control mice that did not experience pre- and post-task sleep sessions. First, we compared the Z-scored velocity isolated from the first half of the reward zone between the two groups during each *reward delay*. Velocity did not significantly vary across groups (F_(1, 36)_ = 1.27, *p* = 0.29), nor did it vary across delays (F_(4, 36)_ = 2.51, *p* = 0.06). Also, there was no interaction present between delay and group (F_(4, 36)_ = 0.30, *p* = 0.88). Although no significant difference was observed in the velocity in the first half of the reward zone, we also evaluated the velocity profile along the entire track during the two *reward delays* (*1 and 1.5 s delays*), where group differences were previously observed in Experiment 1. For the *1 s delay*, velocity varied across the length of the track (F_(59, 531)_ = 1.43, *p* < 0.05), but did not vary significantly across groups (F_(1,531)_ = 0.01, *p* = 0.94), and there was no interaction between group and position on the track (F_(59, 531)_ = 0.65, *p* = 0.98). For the *1.5 s delay*, velocity again varied across the length of the track (F_(54, 486)_ = 1.73, *p* < 0.01). However, velocity did not vary across groups (F_(1,486)_ = 0.18, *p* = 0.68), and there was no interaction between group and position on the track (F_(54, 486)_ = 0.66, *p* = 0.97). Overall, this suggests while 3xTg-AD mice did benefit from sleep, Non-Tg controls did not.

### 3.6 Pathology is not reduced in female 6-month 3xTg-AD mice with sleep sessions

The age at perfusion for the two groups of 3xTg-AD 6-month female mice (**Fig. 6a**; sleep vs. no sleep; t_(12)_ = 1.11, *p* = 0.29) showed that few, if any, extracellular plaques were present at this age. Therefore, we quantified the proportion of cells that were positive for intracellular accumulation of amyloid or pTau, measures we have previously shown to be associated with behavioral performance on this task (Stimmell et al., 2021). We quantified pathology in the regions previously associated with task performance and their importance in spatial navigation, as well as in several control regions: the CA1 field of the hippocampus, parietal cortex (PC), and retrosplenial cortex (RSC) (Clark et al., 2018; Hinman et al., 2018). The dorsal region of the CA1 field (CA1d) was counted separately from the ventral region (CA1v) because the dorsal region is thought to be critical for precise spatial navigation (de Hoz et al., 2003; Moser et al., 1993, 1995; Pothuizen et al., 2004)

**Figure 6.**
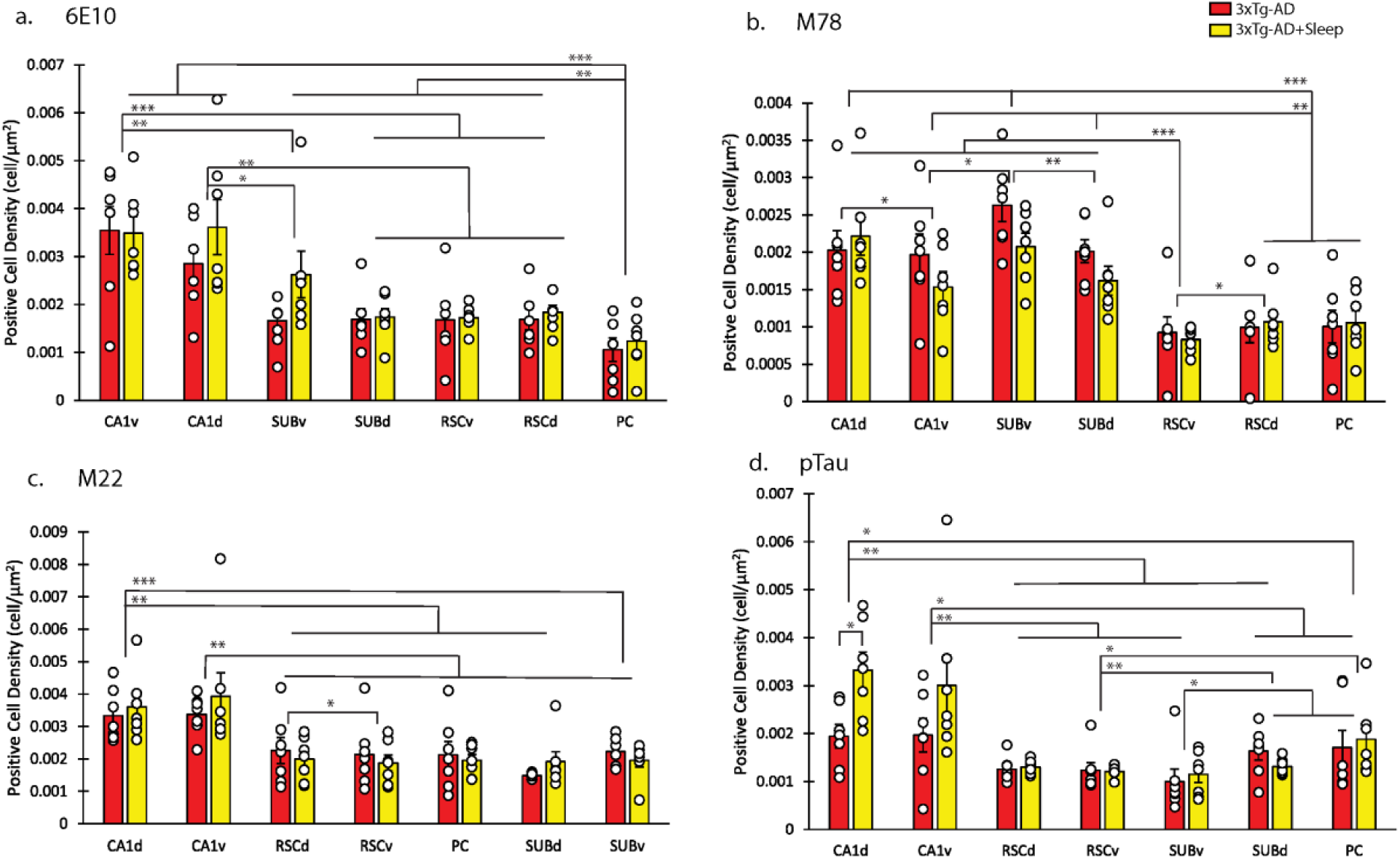
Histological markers of AD pathology do not show a significant decrease in various brain regions after sleep. **a.** Aβ 1-16 specific antibody. **b.** Amyloid fibril conformation specific antibody (mOC78). **c.** Amyloid fibril specific antibody (mOC22). **d.** Antibody specific to hyperphosphorylated isoforms of tau protein (AT8). There was no effect of sleep on Aβ or pTau (*ps* > 0.12). For all stains the group differences are shown above (* p<0.05, ** p<0.01, *** p<0.001).

#### 3.6.1 6E10

There was an effect of brain region (**Fig. 6a**; F_(6, 72)_ = 18.08, p < 0.0001) but no brain region by group interaction (F_(6, 72)_ = 0.94, p = 0.48), and no main effect of group (3xTg-AD and 3xTg-AD+Sleep from *Experiment 1*; F_(1, 12)_ = 1.33, p = 0.27). Planned comparisons for each pair of brain regions to elucidate the source of the variation across brain regions showed that regardless of group, dorsal CA1 (CA1d) and ventral CA1 (CA1v) fields of the hippocampus did not have significantly different densities of intracellular pathology (F_(1, 13)_ = 0.57, *p* = 0.46). Similarly, the dorsal and ventral regions of the subiculum (SUBd and SUBv) were not significantly different in positive cell density (F_(1, 13)_ = 1.90, *p* = 0.19). The dorsal and ventral regions of retrosplenial cortex (RSCd and RSCv) were also not different from each other (F_(1, 13)_ = 0.59, *p* = 0.46). Finally, subiculum (SUB) and retrosplenial cortex (RSC) also were not significantly different (Fs_(1, 13)_ < 1.73, *ps* > 0.21). However, CA1v and CA1d showed significantly higher densities of pathology than all other regions (RSC, SUB, and PC: Fs_(1, 13)_ > 8.76, *ps* < 0.01), while PC had a lower in density than all other regions (CA1, RSC, and SUB: Fs_(1, 13)_ > 10.06, *ps* < 0.01).

#### 3.6.2 M78

There was again an effect of brain region (**Fig. 6b**; F_(6, 72)_ = 24.49, *p*<0.0001) but no brain region by group interaction (F_(6, 72)_ = 1.51, *p* = 0.19), and no main effect of group (F_(1, 12)_ = 0.82, *p* = 0.38). Planned comparisons showed that regardless of group, PC and RSC were not significantly different in intracellular positive cell density (Fs_(1, 13)_ < 1.69×10^-7^, *ps* > 0.10). CA1d was not significantly different from SUB (Fs_(1, 13)_ < 1.86, *ps* > 0.20). CA1v and SUBd were also not significantly different (F_(1, 13)_ = 0.11, *p* = 0.75). However, CA1d showed significantly higher positive cell density than CA1v (F_(1, 13)_=6.02, *p* < 0.05). Similarly, RSCd showed higher positive cell density than RSCv (F_(1, 13)_ = 6.50, *p* < 0.05) but SUBv had higher positive cell density than SUBd and CA1v (Fs_(1, 13)_ > 7.60, *ps* < 0.05). PC and RSC intracellular positive cell densities were both significantly lower than CA1 and SUB (Fs_(1, 13)_ > 18.20, *ps* < 0.01). Interestingly, the conformation specific M78 staining revealed differences in dorsal versus ventral amyloid density that were not apparent with 6E10 (Pensalfini et al., 2014).

#### 3.6.3 M22

We again found an effect of brain region (**Fig. 6c**; F_(6, 72)_ = 15.26, *p* < 0.0001) but no brain region by group interaction (F_(6, 72)_ = 0.84, *p* = 0.54), and no effect of group (F_(1, 12)_ = 0.02, *p* = 0.90). Planned comparisons showed that regardless of group, CA1v and CA1d were not different in intracellular pathology, but both had more than all other regions (RSC, PC, SUB: Fs_(1, 13)_ > 12.89, *ps <* 0.01). RSCd had higher positive cell density than RSCv (F_(1, 13)_ = 10.24, *p <* 0.01). There were no significant differences between SUB, RSC, and PC (Fs_(1, 13)_ < 3.48, *ps* > 0.08). Thus, intracellular amyloid varies across brain regions and the pattern of significant effects varied across the type of amyloid stain, but amyloid is not reduced by sleep.

#### 3.6.4 pTau

We again found an effect of brain region (**Fig. 6d**; F_(6, 72)_ = 11.51, *p* < 0.0001) and for pTau, there was also a brain region by group interaction (F_(6, 72)_ = 2.79, *p* < 0.05), but no main effect of group (F_(1, 12)_ = 2.82, *p* = 0.12). Planned comparison to investigate the significant brain region by group interaction showed that the sleep group had *significantly higher* intracellular positive cell density than the group that did not have rest sessions for CA1d (t_(12)_ = -3.04, *p* < 0.05). When comparing the other groups for each region, no other significant differences were found. When comparing brain regions, regardless of group, to investigate the source of the group main effect, planned comparisons showed that RSCv and SUBv both had significantly less positive cell densities than SUBd and PC (Fs_(1, 13)_ > 4.70, *ps* < 0.04). There was no significant difference between CA1d and CA1v (F_(1, 13)_ = 0.29, *p* = 0.60) which were both higher in positive cell density than all other regions (Fs_(1, 13)_ > 5.40, *ps* < 0.04). No other regions had significant differences in intracellular positive cell density when compared (Fs_(1, 13)_ < 4.65, *ps* > 0.05). Thus, sleep sessions do not reduce intracellular pTau density, which again varied across brain regions. Similar to amyloid stains, the positive cell density tended to be highest in the hippocampus.

## 4. Discussion

We found that the impairment in spatial reorientation learning and memory previously identified (and replicated in this study) in 6-month female 3xTg-AD mice is reversed when daily 50 min sleep sessions are positioned immediately before and after the task (Stimmell et al., 2019).. To confirm that actual sleep, rather than rest, may explain the lack of a spatial reorientation learning and memory impairment in 3xTg-AD mice, we assessed stillness and SWS during the pre- and post-task sleep sessions in recording array mice. We found that mice spent nearly all of these still bouts in either SWS or REM sleep. No significant differences were observed in the proportion of sleep spent still, the proportion of stillness that was spent in SWS, or the average SWS episode length between 3xTg-AD and non-Tg mice, suggesting that differences in sleep characteristics in 3xTg-AD mice are not likely to explain the sleep benefit in these mice. Further, the sleep benefit could be due to enhanced memory consolidation during sleep (Diekelmann and Born, 2010; Gais et al., 2002; Plihal and Born, 1997) or due to the clearance of pathology which occurs during sleep (Buccellato et al., 2022; Verheggen et al., 2018; Xie et al., 2013). Improved cognition could be a result of improving sleep alone (Leinenga et al., 2024). Thus, we assessed Aβ and tau aggregation in sleep versus no-sleep 3xTg-AD mice and found no differences in Aβ or tau aggregation except that there was significantly more tau in the dorsal CA1 fields of the hippocampus of the 3xTg-AD sleep group. Taken together, our data suggest that sleep may be capable of reversing cognitive impairments that occur as a consequence of tau and Aβ aggregation by enhancing memory consolidation rather than altering tau or Aβ levels directly. We ran these experiments in the light cycle to make the conditions more likely to induce sleep, but this may limit translational potential of our findings. Replicating these experiments in humans during the light phase would be important in understanding the effects.

Previous studies have shown that undergoing a sleep session after learning enhances the retention of declarative information, as well as improve performance of procedural skills (Diekelmann and Born, 2010; Gais et al., 2002; Plihal and Born, 1997; Rasch et al., 2007). SWS, specifically, has been associated with facilitating the consolidation of declarative memories (Diekelmann and Born, 2010; Peigneux et al., 2001; Plihal and Born, 1997; Stickgold, 2005). Further, post-learning sleep deprivation has been shown to lead to impaired memory in both humans and animals (Blissitt, 2001; McCoy et al., 2013; Peigneux et al., 2001; Rasch et al., 2007; Williams et al., 1966). Hippocampal-cortical interactions during similar sleep sessions have been shown to be important for memory consolidation and navigation (Cushing et al., 2024, 2020; Jones et al., 2019; Maingret et al., 2016; Oess et al., 2017; Stickgold, 2005; Wang and Morris, 2010). Impaired hippocampal-cortical interactions in 3xTg-AD mice may underlie difficulties in learning a virtual spatial reorientation task (Cushing et al., 2020). Pattern reactivation, which occurs during sleep, may be crucial for memory consolidation, including spatial memories (Ferrara et al., 2008). For example, in rats, hippocampal neuronal activity observed during spatial tasks were shown to re-activate during subsequent sleep in the same spatio-temporal pattern (Born and Wilhelm, 2012; Diekelmann and Born, 2010; Ji and Wilson, 2007; Wilson and McNaughton, 1994). Pattern reactivation has primarily been observed during SWS (Born and Wilhelm, 2012; Creery et al., 2015). There is also evidence for pattern reactivation in humans (Creery et al., 2015). Human neuroimaging studies have shown a similar phenomenon where hippocampal reactivation during sleep was shown to be similar to activity present during prior learning (Bendor and Wilson, 2012; Born and Wilhelm, 2012). During SWS, hippocampal activity in humans was similar to activity present during a prior spatial navigation task (Creery et al., 2015). Interestingly, the 6-month female control mice that underwent pre- and post-task sleep sessions were shown to perform similarly to 6-month female control mice that did not undergo sleep sessions. Thus, the additional sleep may not be as beneficial for control animals.

AD patients experience disrupted sleep, which includes increased wakefulness after the onset of sleep, as well as increased latency to sleep (Peter-Derex et al., 2015). These sleep disorders can be present early on in disease progression and worsen over time (Peter-Derex et al., 2015). Patients experiencing more SWS in post-learning sleep have been shown to perform better on recall of episodic memories (Rauchs et al., 2013). People with AD have been shown to spend less time in SWS and animal models of AD, including the 3xTg-AD mouse model, have been shown to have impaired SWS, even early on in disease progression (Lee et al., 2020). However, impaired sleep in mice modeling tau and/or amyloid aggregation aspects of human AD typically occurs at later timepoints than those examined here, for example, after plaques have begun to form. Interestingly, 6-month female 3xTg-AD and non-Tg mice did not show a difference in the proportion of sleep sessions spent still, the amount of SWS occurring while still, nor the average SWS episode length. Potentially, this could be because the sleep sessions these mice underwent were short rest sessions and not 24 hr sleep. In fact, one month later, while performing a virtual version of this task, these same mice had increased time spent still and increased SWS bout length (Cushing et al., 2020). Given that 6-month female 3xTg-AD sleep mice were not impaired on the spatial reorientation task, sleep, while short, may still be benefiting 3xTg-AD mice by assisting with the consolidation of spatial learning and memory. Interestingly though, 3xTg-AD non-recording array sleep mice spent significantly less time still during their sleep sessions than both non-Tg and 3xTg-AD recording array mice, suggesting 3xTg-AD non-recording array sleep mice spent less time overall in SWS during these sessions. This smaller amount of sleep could potentially be due to having a smaller and lighter implant. However, the 3xTg-AD non-recording array sleep mice are still not impaired at spatial reorientation learning and memory, suggesting that even a smaller amount of sleep can still be beneficial and, while there may be an important threshold for the amount of sleep needed to restore cognition, these mice were able to gain enough to benefit behavior and restore performance to the level of controls. What may be more important is when this sleep is occurring, such as having sleep immediately after a behavioral task, which has been shown to be beneficial in rodents (Diekelmann and Born, 2010; Gais et al., 2002; Plihal and Born, 1997; Rasch et al., 2007).

Sleep has also been linked to removing waste including amyloid and tau aggregates. Conversely, decreased sleep has been shown to lead to a decrease in the clearance of both Aβ and tau, as well as an increase in phosphorylated tau (Lucey, 2020). Both the glymphatic system and the blood-brain barrier (BBB) are thought to participate in the clearance of waste from the interstitial space of the brain (Verheggen et al., 2018). During sleep, the volume of the interstitial space increases, and decreases again upon waking (Xie et al., 2013). Changes in volume here influence the influx of cerebrospinal fluid (CSF), and the exchange with interstitial fluid (ISF) is enhanced (Verheggen et al., 2018). Accumulation of Aβ happens early in disease progression and once enough has accumulated, insoluble plaques form. These plaques can contribute to sleeping problems, which in turn will facilitate further Aβ accumulation (Ju et al., 2014). Enhancing sleep quality and the amount of sleep may help with both decreasing the risk of progression to AD and/or slow or halt AD symptom progression by assisting in the clearance of Aβ and tau (Ragsdale et al., 2024). Previously, we showed that 6-month female 3xTg-AD mice are impaired at spatial reorientation and that these mice had a tau pathology profile across the parietal-hippocampal network that was correlated with spatial reorientation behavior, suggesting a role of tau pathology affecting this behavior (Stimmell et al., 2021, 2019). Thus, in the present study, there may be a potential increase in the clearance of these pathological products, potentially during SWS, which could enhance spatial learning and memory. However, this hypothesis was not supported by the results, as we did not find any reductions in any marker of tau or amyloid aggregation in the parietal hippocampal network. Thus, facilitated memory formation during sleep is the most likely explanation for the improved cognition we observed in 3xTg-AD sleep mice.

In summary, we previously demonstrated that 6-month female 3xTg-AD mice exhibit impaired spatial reorientation and that tau pathology in the parietal-hippocampal network is indicative of this behavior. This study shows that introducing pre- and post-task sleep sessions to 3xTg-AD mice can reverse their spatial reorientation learning and memory impairments, bringing their performance back to control-like levels. Interestingly, despite the 6-month female non-recording array 3xTg-AD mice spending less time still during sleep sessions compared to their recording array counterparts, this was enough to reverse the cognitive impairment. This finding suggests that even a limited amount of sleep may benefit spatial reorientation learning and memory, provided a critical threshold of sleep is met or, crucially, if sleep occurs post-task. Additionally, the lack of reduced pathology in 3xTg-AD sleep mice implies that the cognitive benefits of sleep are likely tied to memory consolidation during sleep rather than the clearance of amyloid or tau. These findings differentiate the impact of sleep-related memory consolidation from amyloid/tau clearance, suggesting that extending sleep duration in the early stages of Alzheimer’s disease could simultaneously facilitate memory consolidation and clear pathology, offering dual benefits to alleviate cognitive impairments.

## Notes

**Funding:** This research was supported by grants from FL DOH 20A09, FL DOH 21K12 (Co-PI), NIA K99/R00 AG049090, NIH R01 AG070094, NIH R56 MH133929 to AAW and NIA 1F31AG079619-01 to SDC.

### Competing Interest Statement

The authors have declared no competing interest.

